# Electro-conductive carbon nanofibers containing ferrous sulfate for bone tissue engineering

**DOI:** 10.1101/2020.12.27.424517

**Authors:** Houra Nekounam, Hadi Samadian, Fatemeh Asghari, Reza Faridi Majidi

## Abstract

The application of electroactive scaffolds can be promising for bone tissue engineering applications. In the current paper, we aimed to fabricate an electro-conductive scaffold based on carbon nanofibers (CNFs) containing ferrous sulfate. FeSO_4_·7H_2_O salt with different concentrations 5, 10, and 15 wt%, were blended with polyacrylonitrile (PAN) polymer as the precursor and converted to Fe_2_O3/CNFs nanocomposite by electrospinning and heat treatment. The characterization was conducted using SEM, EDX, XRD, FTIR, and Raman methods. The results showed that the incorporation of Fe salt did not induce an adverse effect on the nanofibers’ morphology. EDX analysis confirmed that the Fe are uniformly dispersed throughout the CNF mat. FTIR spectroscopy showed the interaction of Fe salt with PAN polymer. Raman spectroscopy showed that the incorporation of FeSO4·7H_2_O reduced the ID/IG ratio, indicating more ordered carbon in the synthesized nanocomposite. Electrical resistance measurement depicted that, although the incorporation of ferrous sulfate reduced the electrical conductivity, the conductive is suitable for electrical stimulation. The *in vitro* studies revealed that the prepared nanocomposites were cytocompatible and only negligible toxicity (less than 10%) induced by CNFs/Fe_2_O_3_ fabricated from PAN FeSO_4_·7H_2_O 15%. These results showed that the fabricated nanocomposites could be applied as the bone tissue engineering scaffold.

## 1. Introduction

Bone tissue engineering is an emerging alternative over conventional bone defect treatments, aiming to eliminate their limitations and drawbacks. Tissue engineering comprises various parts, including tissue scaffolds, cells, bioactive molecules, and bioreactors. The tissue scaffolds play a central role in tissue engineering to support cell attachment, growth, proliferation, and differentiation [1, 2]. Moreover, the scaffolds could be implemented as the local drug delivery vehicle to deliver various bioactive molecules. Various types of biomaterials have been evaluated as the bone tissue engineering scaffolds, for instance, hydrogels, 3D printed structures, freeze-dried structures, and nanofibers. Among them, nanofibrous structures have exhibited fabulous performance as the bone tissue engineering scaffolds due to their resemblance to the extracellular matrix (ECM) of bone tissue, high surface to volume ratio, ability to be fabricated from a wide range of materials, and integration with the different manufacturing process [3]. The resemblance of the prepared tissue engineering scaffold to the native bone structure is crucial for developing efficient tissue engineering scaffolds. Accordingly, plenty of studies have been conducted to develop scaffold with the highest similarity to the native bone structure [4].

Bone is an electroactive tissue and different studies showed that electrical stimulation can improve the bone healing process [5]. Therefore, electroactive and electro-conductive scaffolds have gained significant attention in bone regeneration concept [6–9]. These structures can be applied as the substrate to effectively stimulate the attached cells and modulate the functions of cells. In the bone defect site, electro-conductive scaffolds transmit the electrical signal generated in the bone and accelerate the healing process [10, 11]. Nanofibrous electro-conductive scaffolds could interact more efficiently with bone regenerating cells, osteoblast and osteocytes, due to the nanometric morphology and ECM-mimicking morphology. Electrospun carbon nanofibers (CNFs) possess various promising properties beneficial for bone tissue engineering applications, including proper electrical conductivity, mechanical strength, nanoscale diameters, biocompatibility, and chemical inertness [12].

CNFs can be composited with various nanoparticles (NPs) such as bioactive glass, glass ceramics, alumina, Zirconia, and hydroxyapatite (HA) to promote their performance as tissue engineering scaffolds. In another study [12], we observed that CNFs/HA nanocomposite enhanced bone cell growth and regeneration. Various studies reported the positive effects of metallic NPs on bone regeneration. In our previous study [13] we showed that the incorporation of gold NPs enhanced the performance of CNFs. De Santis et al. [14] fabricated a 3D-scaffold based on poly(ε-caprolactone) (PCL) composited with iron-doped HAp (FeHAp) nanoparticles. They reported 36% higher cell growth using the composited scaffold compared to the pure scaffold. In another study, Kim et al. [15] fabricated PCL-based magnetic scaffold containing magnetic NPs and observed a higher mineral induction and cell adhesion/proliferation on the magnetic scaffold. On the way to develop efficient electro-conductive scaffolds based on CNFs, we blended various concertation of FeSO_4_·7H_2_O salt with polyacrylonitrile (PAN) as the precursor to fabricate Fe2O3 NPs-composited CNFs. The prepared nanocomposites were characterized and the results showed beneficial properties of the prepared structures for bone tissue engineering applications.

## 2. Materials and methods

### 2.1. Materials

PAN (MW: 80,000 g/mol) was obtained from Polyacryl Company (Iran). Ferrous sulfate (FeSO_4·_7H_2_O) were purchased from Sigma-Aldrich (St. Louis MO, USA). N,N-dimethylformamide (DMF) was obtained from Merck (Darmstadt, Germany). Fetal Bovine Serum (FBS), Penicillin, Streptomycin, and DMEM/F12 were purchased from Gibco (Gaithersburg, MD, USA). Lactate dehydrogenase (LDH) assay kit (LDH Cytotoxicity Detection KitPLUS) and MTT Kit were purchased from Roche (Manheim, Germany). Human osteosarcoma cells (MG-63) were supplied by the National Cell Bank of Iran (NCBI) based in the Pasteur Institute of Iran (NCBI, C555).

### 2.2. Electrospinning

PAN was dissolved in DMF with a concentration of 9% w/v (0.09 g/mL) and stirred at 40 °C for 12 h to obtain a homogenous and clear solution. Different concentrations of FeSO_4_·7H_2_O, 5, 10, and 15 wt.%, were added to the prepared PAN solution and stirred at 60 °C for 24 h to obtain a heterogeneous solution. A commercial electrospinning system (Fanavaran Nano Meghyas Ltd., Co., Tehran, Iran) was used to make nanofibers from the prepared solutions. The electrospinning parameters were as follows the applied voltage of 20 kV, feeding rate of 1 mL/h, and nozzle to collector distance of 10 cm. The resulted nanofibers were converted to CNFs via two steps heat treatment in a tube furnace (Azar, TF5/25– 1720, Iran); stabilization at 290 °C for 3 h with a heating rate of 1.5 °C/min and carbonization at 1000 °C for 1h under a high-purity nitrogen atmosphere (N2 99.9999 %, Air Products) with the heating rate of 4 °C/min.

### 2.3. Characterization of nanofibers

The prepared nanofibers were characterized before and after carbonization. Scanning Electron Microscopy (SEM) equipped with Energy Dispersive X-Ray (EDX) detector (Philips XL-30, at 20 kV) was used to observe the microstructure and morphology of the nanofibers and assess the semi-quantitative elemental analysis. Nanofibers were sputter-coated with a thin layer of gold using a sputter coater (SCD 004, Balzers, Germany) and observed using SEM. The Image J software (1.47v, National Institute of Health, USA) was used to calculate the fibers diameter.

Fourier transform infrared (FTIR) spectrum of nanofibers were recorded using a Shimadzu 8101M FTIR (Kyoto, Japan) at room temperature. The crystallinity of the prepared nanofibers was evaluated by recording the XRD pattern using Siemens D5000 diffractometer (Aubrey, TX, USA) and X-ray generator (Cu Kα radiation with λ= 1.5406 Å, 40 kV, 30 mA, the step size of 0.08°/s range from 2° to 80° (2θ)) under ambient conditions. Raman spectra of the nanofibers was recorded using a SENTERRA spectrometer (BRUKER, Germany, 785 nm, 25 mW, resolution of ~3.0 cm−1 over a range of 90–3500 cm−1) to assess the homogeneous nature of the samples. The nanofibers’ wettability was investigated based on the water contact angle (WCA) method using a static contact angle measuring device (KRUSS, Hamburg, Germany). The electrical resistivity of the prepared nanofibers was measured using a 4-point probe multi-meter (Signatone SYS-301 with Keithley 196 system DDM multimeter) and the conductivity was worked out based on Equation 1.

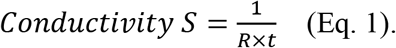

Where R and t are the sheet resistance in ohms and the film thickness in cm, respectively.

### 2.4. Cytotoxicity and proliferation evaluations

The toxicity of the fabricated nanocomposite on MG-63 cells was evaluated by direct and indirect methods using LDH and MTT assay kits, respectively. For direct toxicity measurement, nanocomposites were cut circulatory, located at the bottom of the 96-well plate. Sterilization was conducted using ethanol (70.0%) for 2 h and UV light irradiation for 20 min and washing thoroughly with sterile PBS (pH 7.4). A number of 5 × 10^3^ cells suspended in 100 μL of culture medium supplemented with FBS (1.0% v/v) and antibiotic (1.0%) were seeded on the nanocomposites and incubated at 37 °C in a humidified atmosphere. The toxicity induced by nanocomposites was measured 24, 48, and 72 h after cell seeding based on Equation 2.

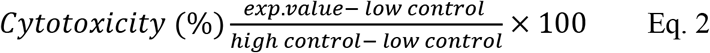

Low control was LDH released from untreated cells and high control was the LDH content of cells obtained by lysing the cells. The toxicity percent was relative to the negative control, cells seeded on the tissue culture plastics (TCP).

The indirect toxicity was conducted according to our previous study [5] and based on the ISO 10993-5 standards. Briefly, 0.1 g of nanocomposites were incubated in 1 mL of cell culture medium containing 1.0% antibiotic and 10% FBS for 7 days at 37 °C. The same amount of cultured medium without nanocomposites was incubated in the same conduction and considered as the control. The incubated culture mediums were extracted and restored for the indirect toxicity assay. A number of 5 × 10^3^ cells suspended in 100 μL of culture medium supplemented with FBS (1.0% v/v) and antibiotic (1.0%) were seeded in the 96-well plate and incubated at 37 °C in a humidified atmosphere for 24 h. Then, 100 μL of the extracted culture medium was replaced with cell culture medium and incubated for another 24 h. After the incubation time, the cell culture medium was aspirated and replaced with MTT solution (0.5 mg/ml) and incubated for 4 h. Finally, the formed formazan crystals were dissolved by DMSO and the absorbance was measured using a microplate reader (Anthos 2020, Biochrom, Berlin, Germany) at 570 nm.

#### 2.4.1. Proliferation measurement

The proliferation of MG-63 cells on the prepared nanocomposites was measured using LDH assay kit. The nanocomposites were cute circulatory, put in 96-wells tissue culture plate, sterilized, and seeded with the cells according to the methods described before. Cell proliferation was measured at 24, 48, and 72 h post-cell seeding according to the total LDH amount of cells [12]. After the time points the cells lysed with the lysing solution and the LDH amount measured at 490 nm using the microplate reader (Anthos 2020, Biochrom, Berlin, Germany). Equation 3 Was used to calculate the proliferation of cells.

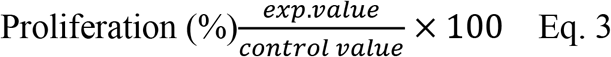

### 2.5. Cells morphology observation

The morphology of MG-63 cells on the prepared nanocomposites was observed using SEM. The cells were fixed with paraformaldehyde (4% w/w) for 1 h and dehydrated in upgrading ethanol. The fixed cells were putter coated with a thin layer of gold and observed using the SEM apparatus at an accelerating voltage of 26 kV.

### 2.6. Statistical analysis

The experiments were performed triplicate, except for the *in vitro* cell culture assays which repeated five times. The obtained results were statistically analyzed using SPSS (version 10.0) by One-way ANOVA (ANalysis of VAriance). The results were reported as a mean ± standard deviation (SD) and p < 0.05 set as the significance level.

## 3. Results and discussion

### 3.1. Characterization results

The prepared nanocomposites were characterized by different methods. The nanofibers’ morphology was observed by SEM, and the presence of Fe in the nanofibers confirmed by the EDX method (Figs. 1 and 2).

**Fig. 1.**
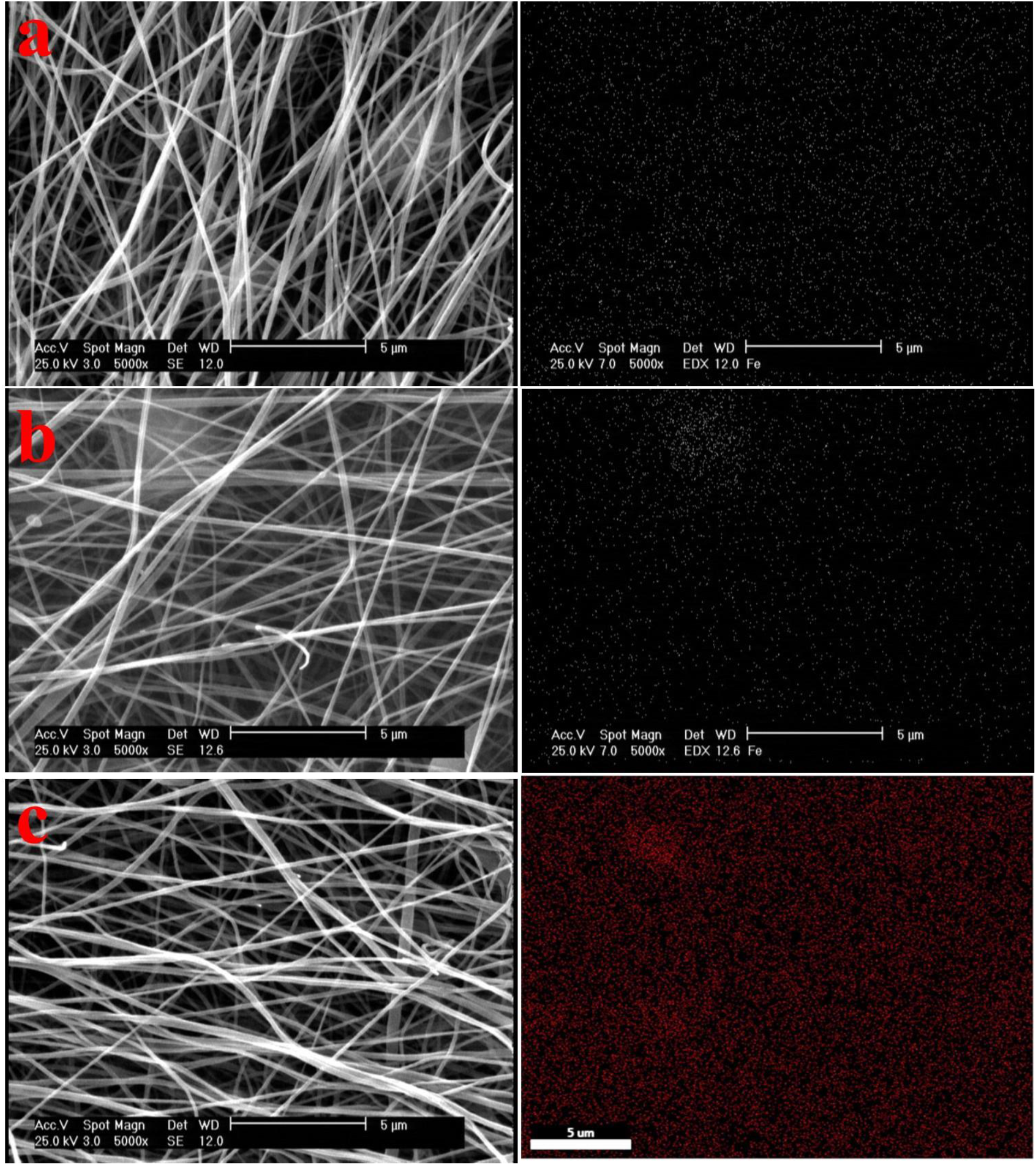
SEM micrograph and EDX map of FeSO_4_·7H_2_O-PAN nanofibers with different concentrations of FeSO_4_·7H_2_O. a) 5, b) 10, and c) 15 wt.%.

**Fig. 2.**
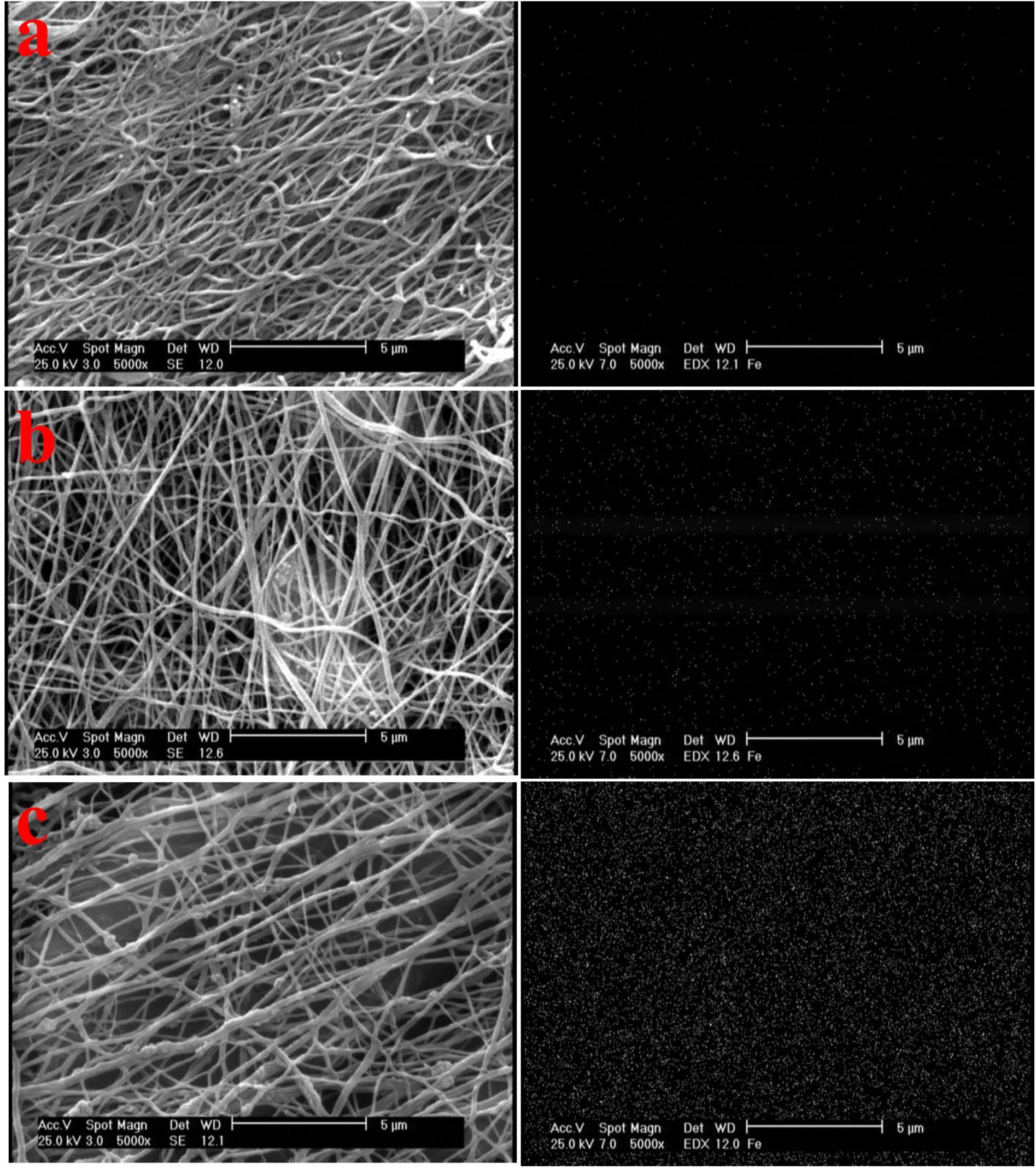
SEM micrograph and EDX map of CNFs prepared from FeSO_4_·7H_2_O-PAN nanofibers with different concentrations of FeSO_4_·7H_2_O. a) 5, b) 10, and c) 15 wt.%.

The SEM images showed that the incorporation of the ferrous sulfate had no adverse effects on the morphology of the PAN nanofibers and the fibers are straight, uniform, and beadles. Moreover, it was observed that increasing the concentration of ferrous sulfate decreased the PAN nanofibers diameter [16, 17]. Ji et al. reported the same pattern and showed that increasing the concentrations of FeCl_3_·6H_2_O [16] and Ni(OAc)_2_·4H_2_O [17] resulted in PAN nanofibers with bigger diameter. EDX analysis confirmed the presence of Fe ions in the electrospun PAN nanofibers. The prepared PAN nanofibers were converted to CNFs by heat treatment and morphology observations and EDX analysis results are presented in Fig. 2.

SEM images of the carbonized FeSO_4_·7H_2_O-PAN nanofibers show some structural deformations and fissions in nanofibers because of PAN polymer chains’ interaction with oxygen and formation of ladder-like structure [5]. Moreover, the diameter of CNFs was reduced during the carbonization process due to removing various components during the process. The average diameter of CNFs with different FeSO_4_·7H_2_O concentrations of 5, 10, and 15 wt % in PAN nanofibers precursor are proximately 102, 126, and 138 nm, respectively. Moreover, EDX analysis confirmed the presence of Fe element in CNFs. The concentration-dependent effect of FeSO_4_·7H_2_O on the EDX map of the nanocomposite is apparent in the EDX results. Furthermore, EDX map analysis showed uniform dispersion of Fe onto/into CNF mat.

#### 3.1.1. FTIR spectra

Surface functional groups of the nanofibers and possible interactions between them were evaluated using FTIR spectroscopy and the results are presented in Fig. 3. The carboxylic group peaks of pure CNFs located at 1550 and 1707 Cm^−1^ were disappeared in CNFs/FeNPs nanocomposites, indicating the interaction between Fe salt and PAN polymer. Moreover, the sulfate esters peaks located at 1414 Cm^−1^ and 1383 Cm^−1^ in CNFs/Fe 5% and CNFs/Fe 15%, respectively, and sulfoxide peak located at 1032 Cm^−1^ in CNFs/Fe 10% confirmed the presence of applied salt in the nanocomposites.

**Fig. 3.**
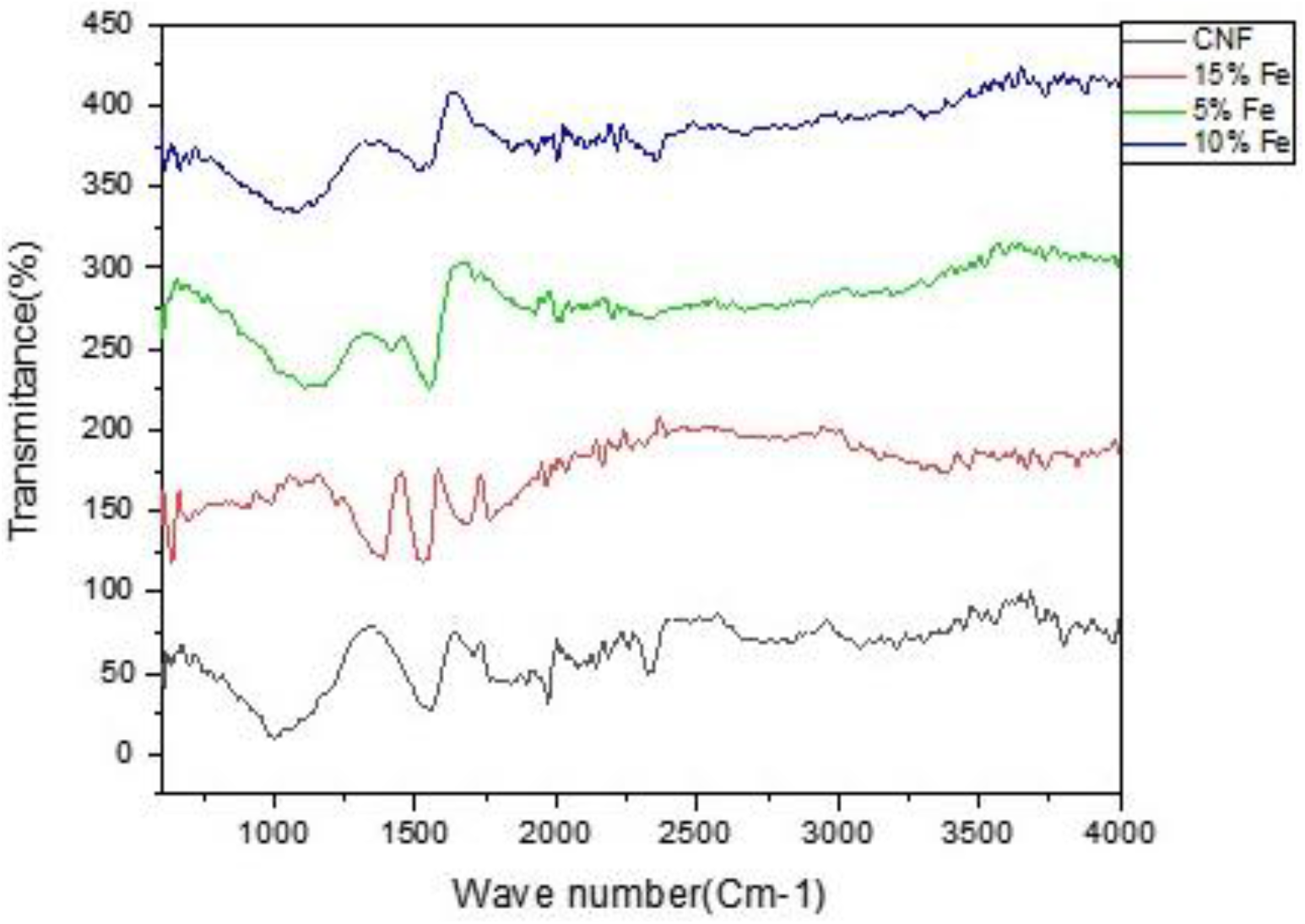
FTIR spectra of CNFs prepared from FeSO_4_·7H_2_O-PAN precursors with different FeSO_4_·7H_2_O concentrations

**Fig. 3.**
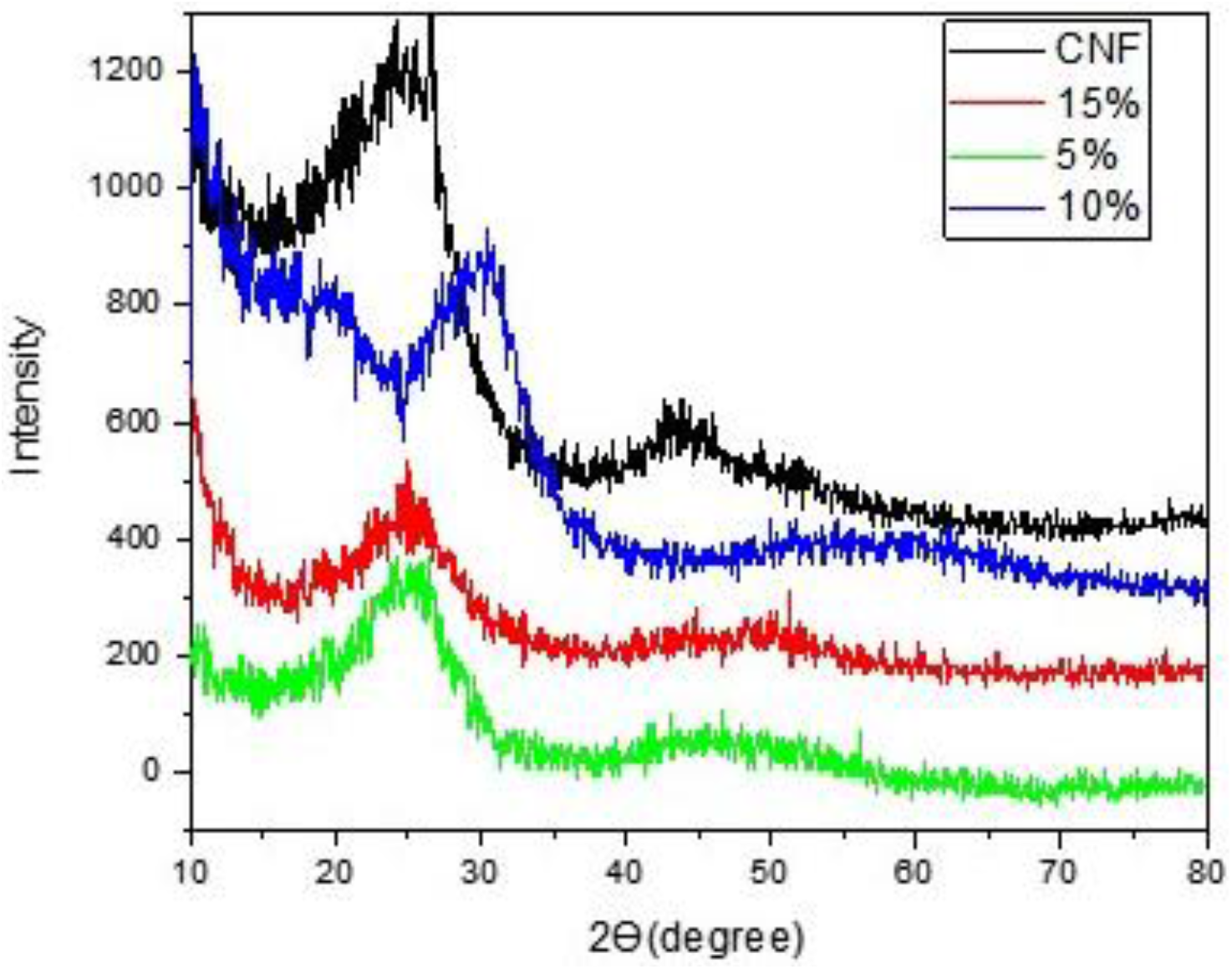
XRD patterns CNFs prepared from FeSO_4_·7H_2_O-PAN precursors with different FeSO_4_·7H_2_O concentrations

#### 3.1.2. XRD pattern

The crystallinity of the prepared nanocomposite was evaluated using XRD, and the results are presented in Fig. 4. The peaks located at 2Ө=26° and 2Ө=44° are corresponding to (001) and (002) planes, respectively, indicating the partial crystallinity of the nanocomposites. The peak located at 2Ө= 35.63 can be related to to the crystallographic plane (104) of hematite with rhombohedral shape and a, b, and c values of 5.03, 5.03, and 13.74, respectively, and α, β, and ɣ values of 90, 90, and 120, respectively.

**Fig. 4.**
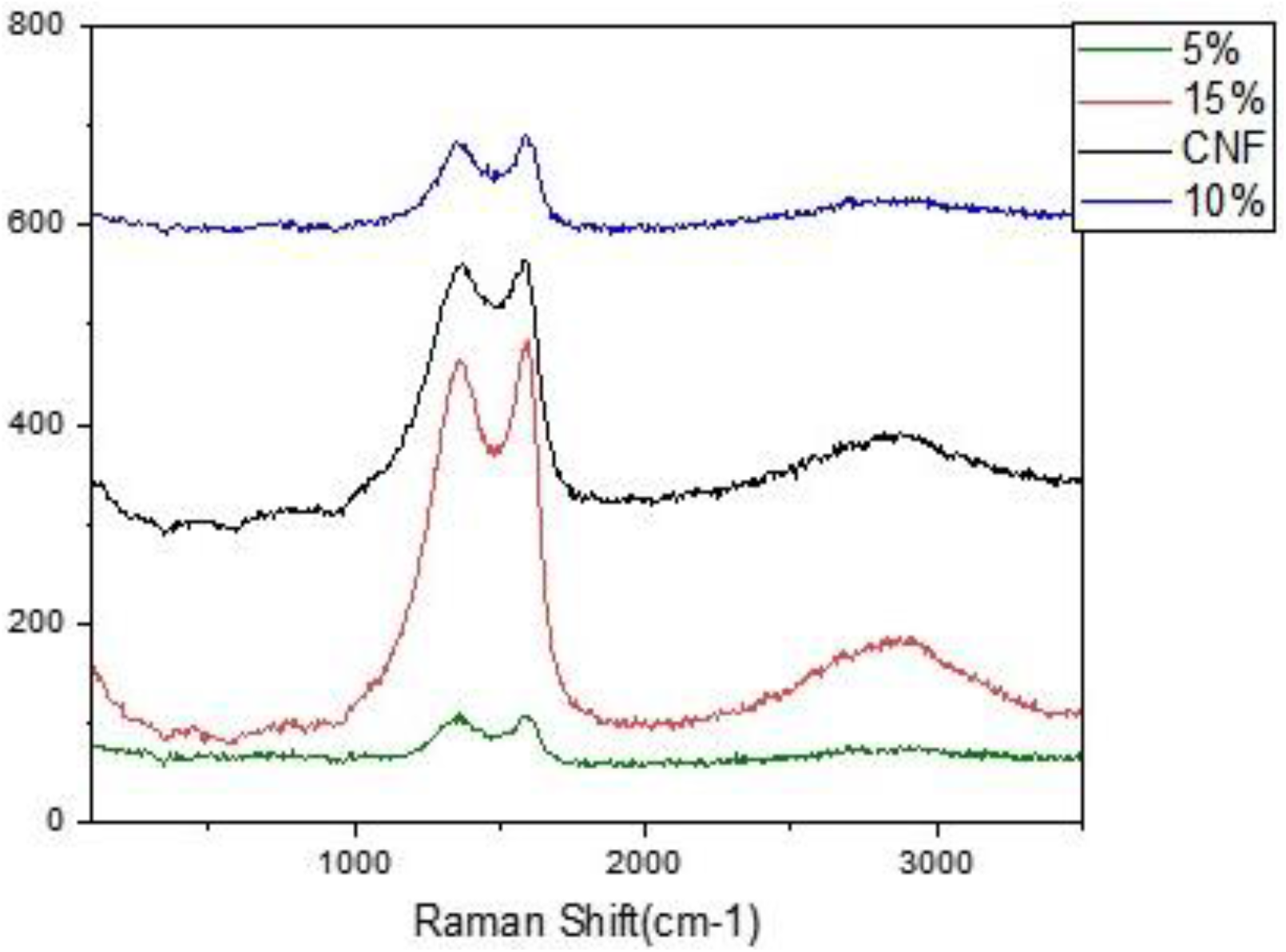
Raman spectra of CNFs prepared from FeSO_4_·7H_2_O-PAN precursors with different FeSO_4_·7H_2_O concentrations

It was observed that at the highest concentration of FeSO_4_·7H_2_O, the hematite crystalline structure reduced the characteristic peak width of CNFs, indicating the reduction of the amorphous nature of CNFs. While at the lower concentrations, the amorphous structure of CNFs dominated the crystallinity of the hematite. Moreover, according to XRD software analysis (Table 1), the incorporation of FeSO_4_·7H_2_O reduced CNFs crystal size and crystalline plane distance.

**Table 1.** Electrical resistivity/conductivity of CNFs prepared from FeSO_4_·7H_2_O-PAN precursors with different FeSO_4_·7H_2_O concentrations.

|  | CNF | 5% Fe | 10% Fe | 15% Fe |
| --- | --- | --- | --- | --- |
| **Conductivity(s.cm<sup>-1</sup>)** | 2.74±0.02 | 1.31±0.14 | 0.98±0.23 | 0.53±0.19 |
| **Resistivity($\Omega$.cm)** | 0.36±0.027 | 0.76±0.06 | 1.02±0.02 | 1.88±0.05 |

#### 3.1.3. Raman spectra

Fig. 4. shows the Raman spectroscopy results of CNFs and nanocomposites with different ferrous sulfate contents. Well-known D-band between 1250 and 1450 cm−1 (disorder-induced phonon mode) and G-band in the range of 1550-1660 cm−1 (graphite band) were observed in all samples. The D-band can be related to disordered and defective portions of carbon (sp3 –coordinated) and the G-band can be attributed to the graphitic crystallites of carbon (sp2 –coordinated) [18–20].

The number of carbon defects in the prepared nanocomposites can be assessed using the relative intensities of the I_D_/I_G_ ratio (Table 2). The lower I_D_/I_G_ ratio represents a large amount of sp2 – coordinated carbon [16, 18]. The results show that the incorporation of FeSO_4_·7H_2_O reduced the I_D_/I_G_ ratio, indicating more ordered carbon in the synthesized nanocomposite.

#### 3.1.4. Water contact angle values

The prepared nanocomposites’ wettability was evaluated using the water contact angle method, and the results are presented in Fig, 5. The results showed that the incorporation of FeSO_4_·7H_2_O had a negligible effect on the wettability of the CNFs. The water contact angle value of the pure CNFs was around 94.2 ° and increased to 108.7, with the highest concentration of FeSO_4_·7H_2_O (15 wt.%).

**Fig. 5.**
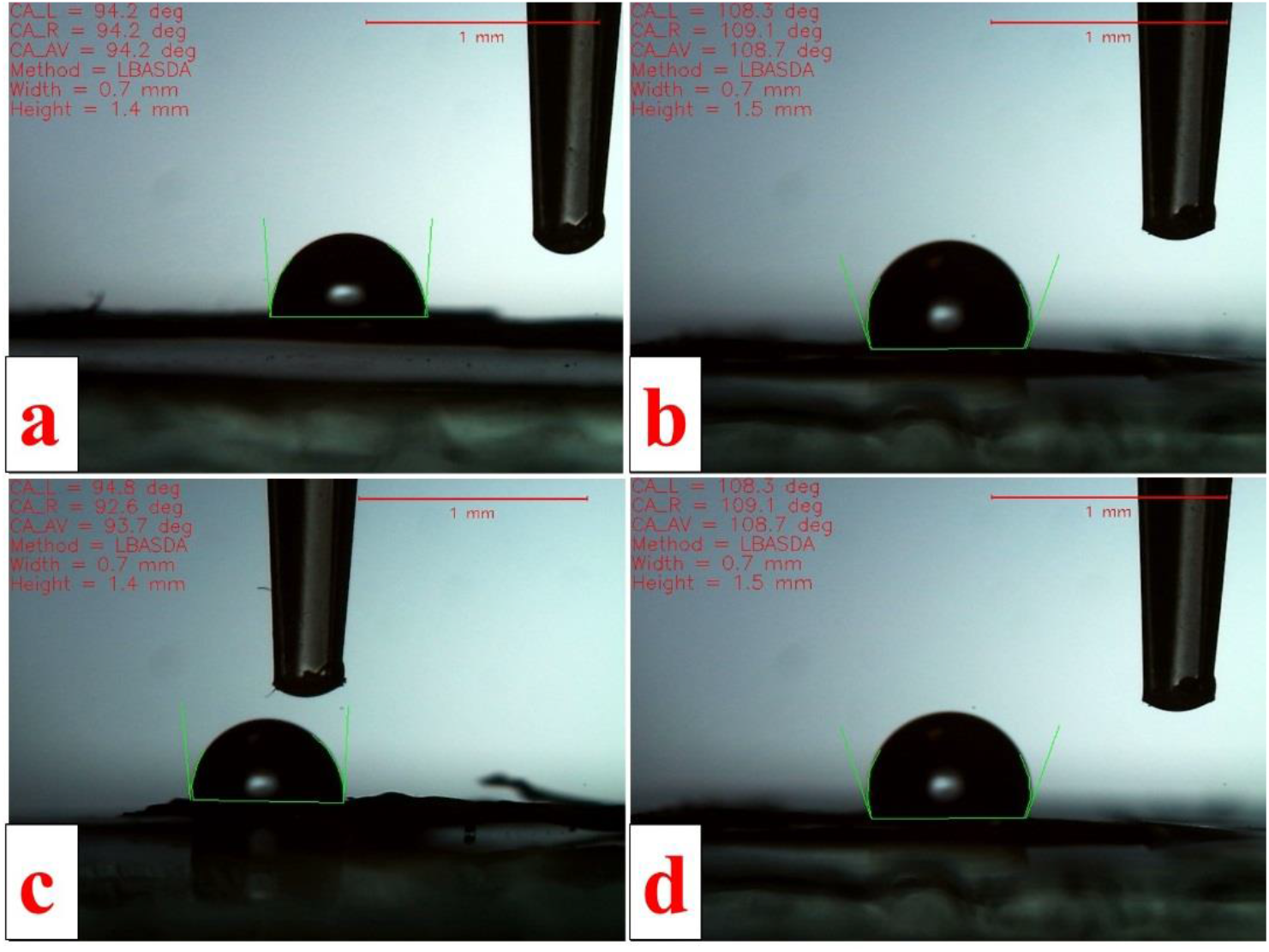
Water contact angle values of CNFs prepared from FeSO_4_·7H_2_O-PAN nanofibers with different concentrations of FeSO_4_·7H_2_O. a) 0, b) 5, c) 10, and d) 15 wt.%

The electrical resistivity of the prepared nanocomposites was measured using a 4-point probe and the results are presented in Table 3. The results showed that the incorporation of the Fe NPs precursor increased the resistivity and reduced the conductivities. These observations could be attributed to the effect of NPs on the consistence of carbon structures in the architecture of the nanocomposite.

#### 3.1.5. Mechanical flexibility

Generally, electrospun CNFs are brittle and their low flexibility hampered their biomedical applications [5]. In the current study, we observed that the prepared pure CNFs were brittle under bending (Fig. 6b) and FeSO_4_·7H_2_O-PAN-drived CNFs were flexible (Fig. 6b). The flexible CNFs mat is more suitable for tissue engineering applications.

**Fig. 6.**
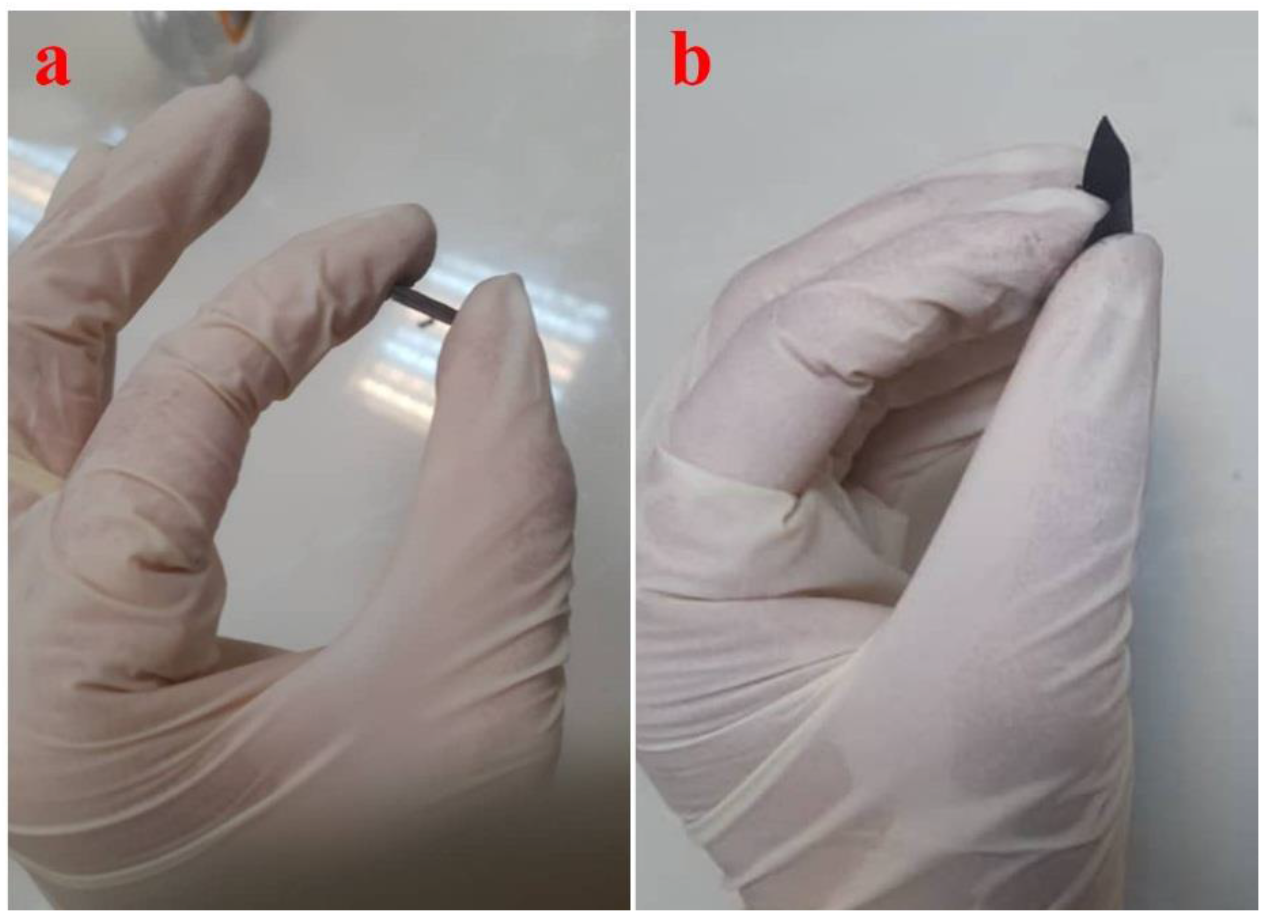
Flexibility of (a) pure CNFs and (b) FeSO_4_·7H_2_O-PAN-drived CNFs under bending force.

### 3.2. *In vitro* evaluations

#### 3.2.1. Direct toxicity results

The prepared nanocomposites’ direct toxicity was measured using the LDH assay kit at 24, 48, and 72 h post cell seeding and, the results are presented in Fig. 7. The results showed that the incorporation of FeSO_4_·7H_2_O did not induce significant adverse effects on cell viability.

**Fig. 7.**
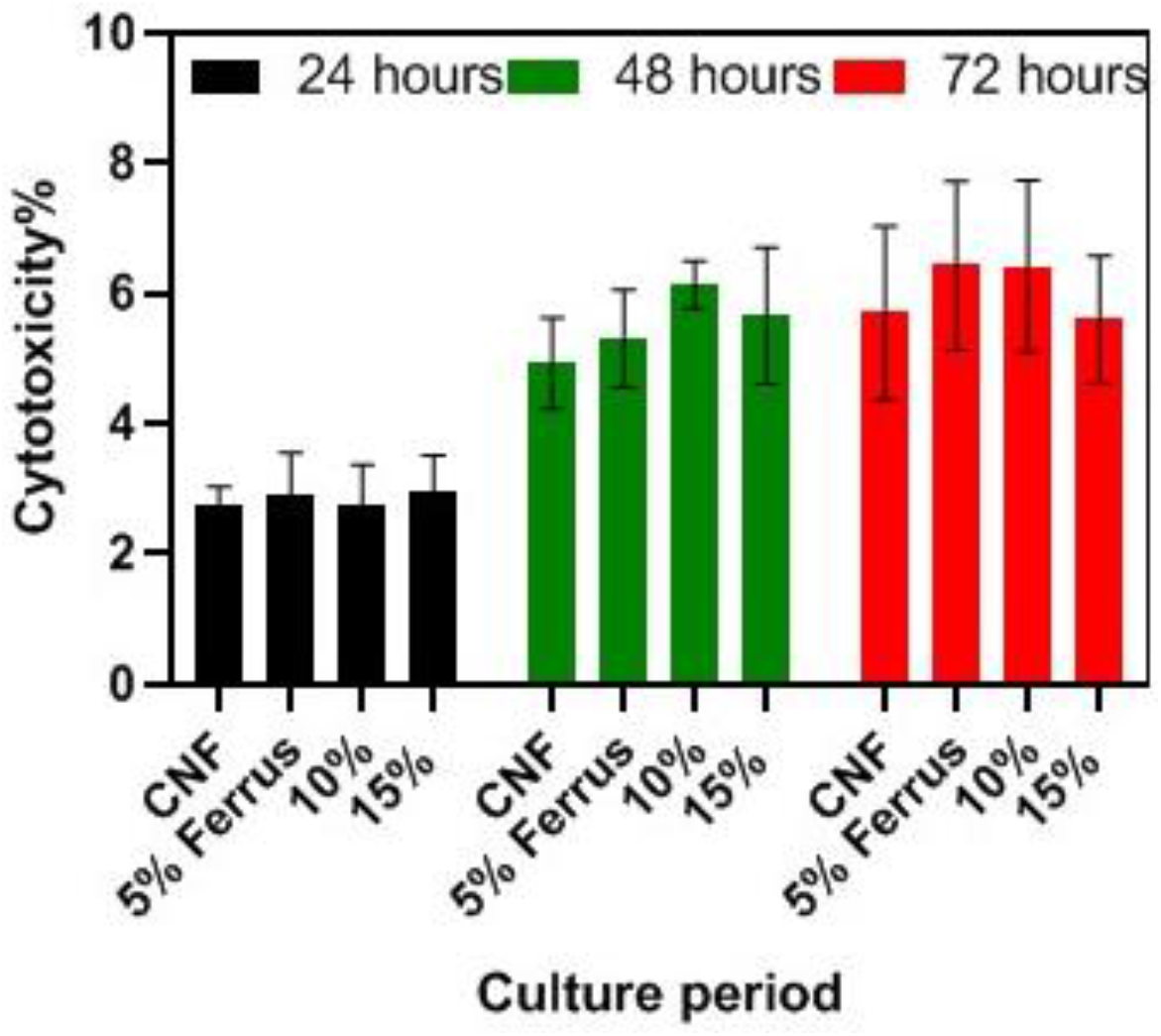
Direct cytotoxic effects of CNFs prepared from FeSO_4_·7H_2_O-PAN nanofibers on MG-63 cells, measured by lactate dehydrogenase (LDH) assay. MG-63 cells seeded on CNFs with a density of 5,000 cells per well in a 96-well plate and incubated for 24, 48, and 72 h. Data represented as mean ± SD, n = 5. The results are given relative values to the negative control (tissue culture plate, TCP).

Moreover, the observed toxicity percent was lower than 10%, suitable for tissue engineering applications.

#### 3.2.2. Indirect toxicity results

The effect of the possible leaked substances from the nanocomposites on the cell viability was measured using the indirect toxicity method. The results showed that the extracts did not induce any adverse effects on the cell viability and the cell viability of the test groups were higher than the control group (Fig. 8). These results confirmed that there are not any leachable substances in the structure of the nanocomposite or the possible leaked substances are not toxic. The toxicity evaluations, direct and indirect methods, implied that nanofibers’ observed toxic effect could be attributed to the structural characteristics rather than leaked substances.

**Fig. 8.**
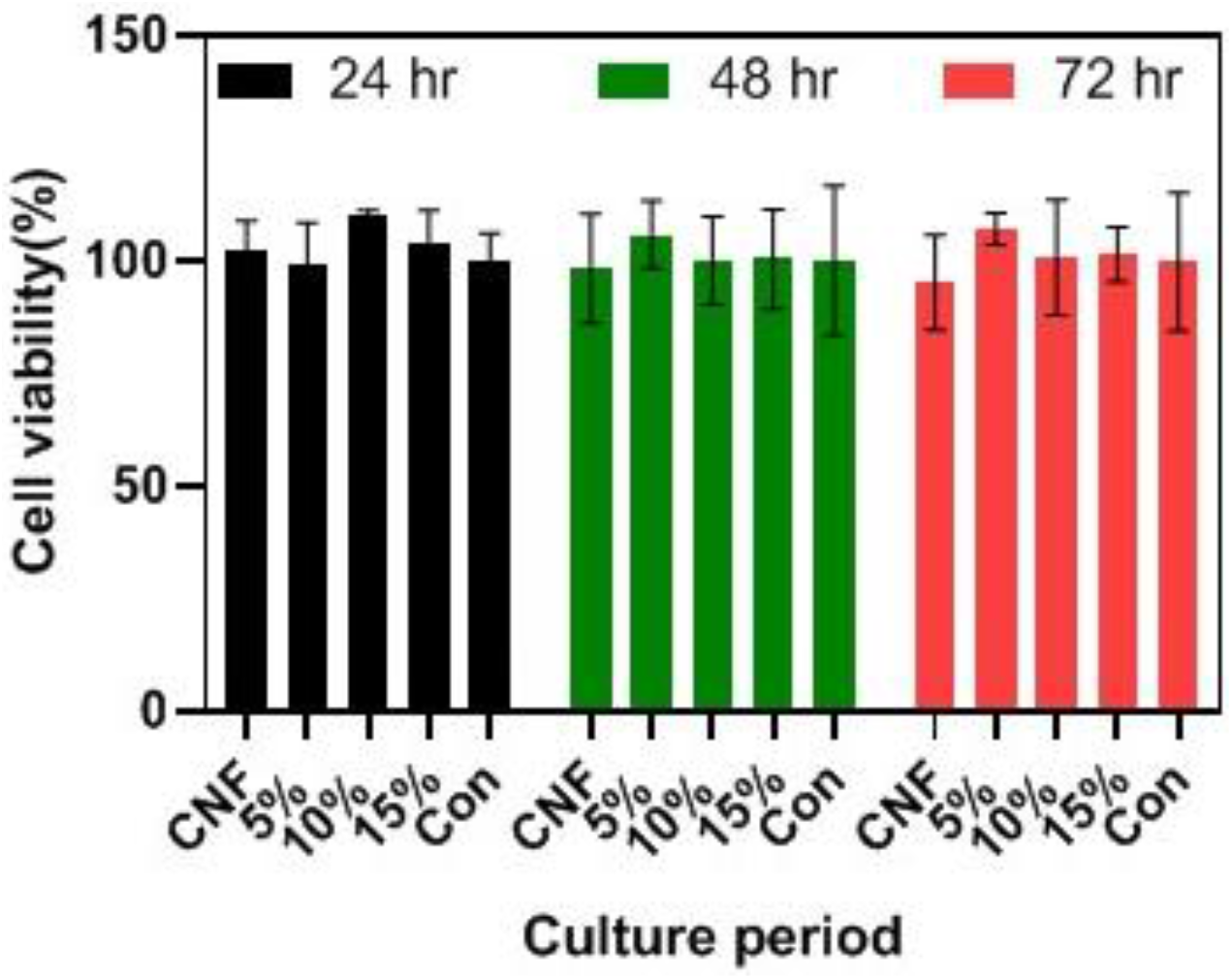
Indirect cytotoxic effects of CNFs prepared from FeSO_4_·7H_2_O-PAN nanofibers on MG-63 cells, measured by MTT assay. MG-63 cells were incubated with the nanofibers extract for 24, 48, and 72 h. Data represented as mean ± SD, n = 5. The results are given relative values to the negative control (tissue culture plate, TCP).

#### 3.2.3. Proliferation results

The proliferation of the cells on the prepared nanocomposites was measured based on the total LDH activity assay, and the results are presented in Fig. 9. The proliferation of MG-63 cells on CNF 5 and 10% insignificantly increased compared to the control and pure CNFs groups (p < 0.05). The proliferation of cells on CNF 15% was lower than the control group indicating the nanofibers’ toxic performance. These results revealed that the prepared nanocomposites are biocompatible and can be applied as the bone tissue engineering scaffolds/ structure.

**Fig. 9.**
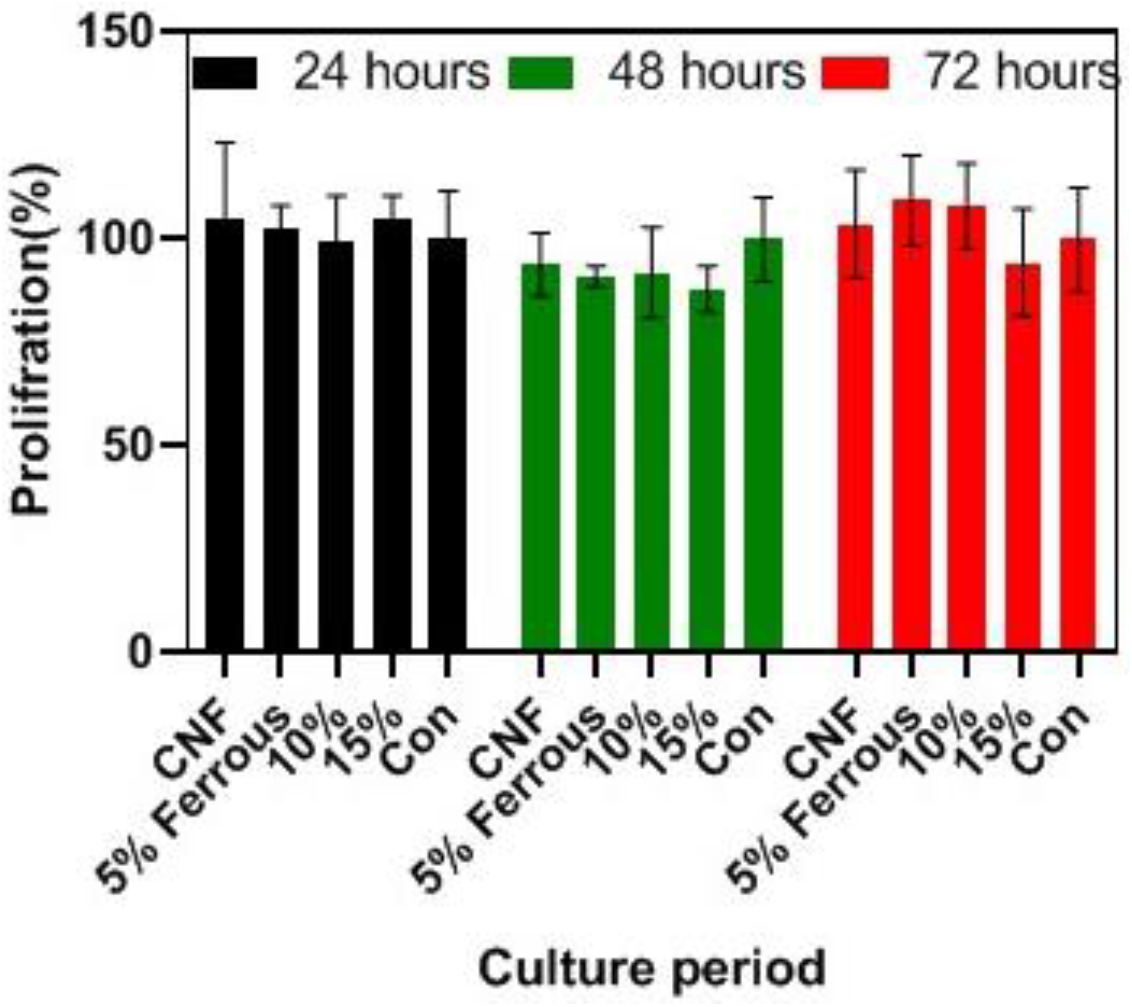
The proliferation of MG-63 cells on CNFs prepared from FeSO_4_·7H_2_O-PAN nanofibers measured by total lactate dehydrogenase (LDH) activity assay. The MG-63 cells seeded on CNFs with a density of 5,000 cells per well in a 96-well plate and incubated for 24, 48, and 72 h. LDH activity was measured after cell lysis. Data represented as mean ± SD, n = 5. The results are given relative values to the control (tissue culture plate, TCP) in percent, whereas the proliferation of cells on control is set to 100%.

#### 3.2.4. Cell attachment and morphology

Cell attachment and maintain its native morphology onto scaffolds are critical for performing their biological activities. The morphology of cells on the nanofibers was observed using SEM imaging and the results are presented in Fig. 10. The results showed that the MG-63 cells are appropriately attached and spread onto the CNF and α-Fe_2_O_3_-CNFs.

**Fig. 10.**
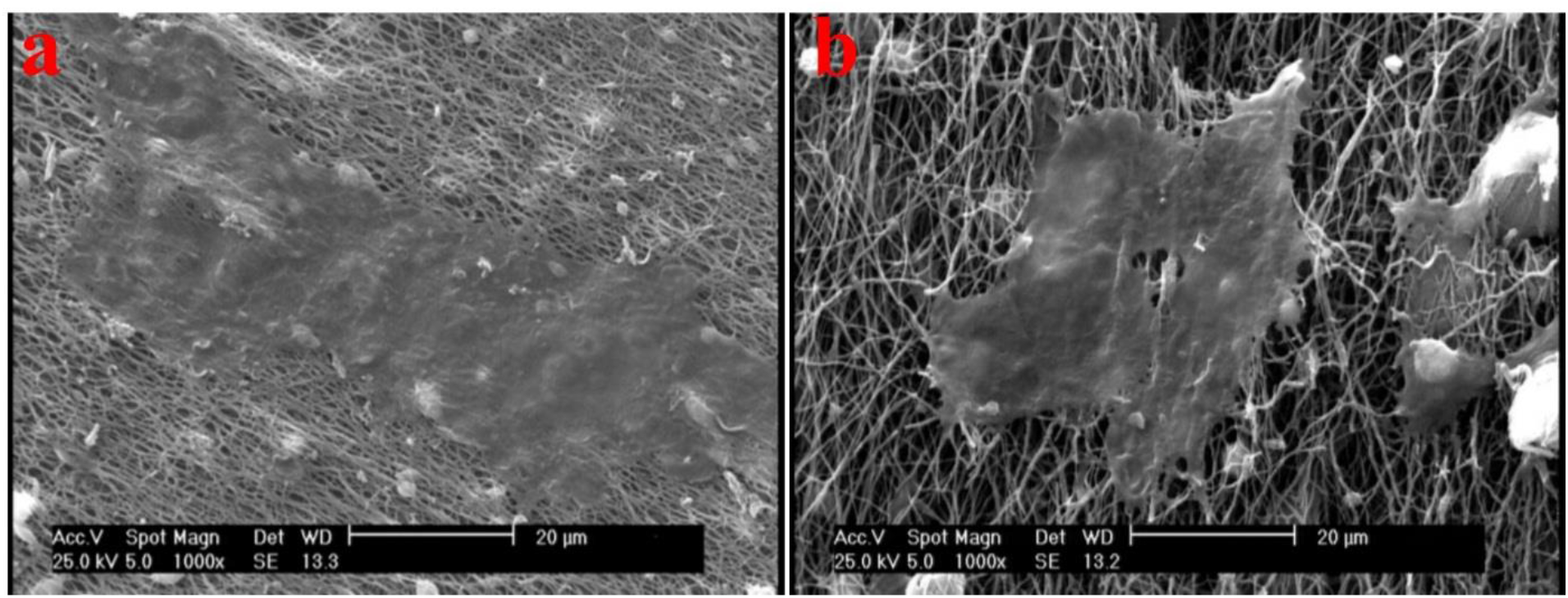
Mg-63 cells attachment and morphology on the prepared nanocomposites. (a) pure CNFs and (b) FeSO_4_·7H_2_O-PAN-drived CNFs

## 4. Conclusion

The development of advanced and promising structures as the bone tissue engineering scaffolds is a breakthrough toward an efficient bone healing approach. Accordingly, in the current study, we fabricated CNFs from PAN nanofibers containing different concentrations of feus sulfate (5, 10, and 15 wt.%) as the precursor nanofibers and evaluated the physicochemical and biological properties of the prepared nanocomposite. SEM images showed that the NPs are embedded into the CNFs and had no adverse effects on the morphology of PAN nanofibers and CNFs. EDX analysis confirmed the presence of Fe and also the uniform dispersion of the FE throughout the nanofibrous mat. The wettability of the prepared nanofibers was slightly reduced by incorporating the NPs, which was not statistically significant (p < 0.2).

FTIR analysis showed the interaction between Fe salt and PAN polymer and also the presence of the applied salt in the nanocomposite. XRD pattern showed that the prepared nanocomposite was a combination of crystalline and amorphous carbon. Raman spectroscopy showed that the incorporation of FeSO4·7H2O reduced the ID/IG ratio, indicating more ordered carbon in the synthesized nanocomposite. It was observed that the electrical resistivity of the nanocomposites increased with the addition of the ferrous salt. The *in vitro* studies showed that the synthesized nanocomposite was biocompatible, and negligible toxicity was observed in CNF/feus sulfate, which was lower than 10%. The MG-63 attachment and morphology on the nanofibers depicted that the cells appropriately spread onto the nanofibers indicating the suitability of the nanofibers for cell adhesion and proliferation. In conclusion, these results showed that CNFs composited with Fe NPs can be applied as the bone tissue engineering scaffolds/structures. It is necessary to evaluate the bone healing efficacy of the prepared nanocomposite in a proper animal model.

## Acknowledgments

We thank our colleagues in Tehran University of Medical Sciences and National Cell Bank of Iran, Pasteur Institute of Iran. This work was supported by Tehran University of Medical Sciences, grant no. 98-01-87-41015.

## References

1. Lin, W., et al., Three-dimensional electrospun nanofibrous scaffolds for bone tissue engineering. Journal of Biomedical Materials Research Part B: Applied Biomaterials, 2020. 108(4): p. 1311–1321.

2. Eivazzadeh-Keihan, R., et al., Metal-based nanoparticles for bone tissue engineering. Journal of Tissue Engineering and Regenerative Medicine, 2020.

3. Samadian, H., et al., Sophisticated polycaprolactone/gelatin nanofibrous nerve guided conduit containing platelet-rich plasma and citicoline for peripheral nerve regeneration: In vitro and in vivo study. International journal of biological macromolecules, 2020. 150: p. 380–388.

4. Koons, G.L., M. Diba, and A.G. Mikos, Materials design for bone-tissue engineering. Nature Reviews Materials, 2020: p. 1–20.

5. Nekounam, H., et al., Electroconductive Scaffolds for Tissue Regeneration: Current opportunities, pitfalls, and potential solutions. 2020: p. 111083.

6. Jaymand, M., et al., Development of novel electrically conductive scaffold based on hyperbranched polyester and polythiophene for tissue engineering applications. Journal of Biomedical Materials Research Part A, 2016. 104(11): p. 2673–2684.

7. Massoumi, B., et al., Electrically conductive nanofibrous scaffold composed of poly (ethylene glycol)-modified polypyrrole and poly (∊-caprolactone) for tissue engineering applications. Materials Science and Engineering: C, 2019. 98: p. 300–310.

8. Hatamzadeh, M., et al., Electrically conductive nanofibrous scaffolds based on poly (ethylene glycol) s-modified polyaniline and poly (∊-caprolactone) for tissue engineering applications. Rsc Advances, 2016. 6(107): p. 105371–105386.

9. Massoumi, B., et al., A novel bio-inspired conductive, biocompatible, and adhesive terpolymer based on polyaniline, polydopamine, and polylactide as scaffolding biomaterial for tissue engineering application. International journal of biological macromolecules, 2020. 147: p. 1174–1184.

10. Gholizadeh, S., et al., Preparation and characterization of novel functionalized multiwalled carbon nanotubes/chitosan/β-Glycerophosphate scaffolds for bone tissue engineering. International journal of biological macromolecules, 2017. 97: p. 365–372.

11. Allahyari, Z., et al., Optimization of electrical stimulation parameters for MG-63 cell proliferation on chitosan/functionalized multiwalled carbon nanotube films. Rsc Advances, 2016. 6(111): p. 109902–109915.

12. Samadian, H., et al., Osteoconductive and electroactive carbon nanofibers/hydroxyapatite nanocomposite tailored for bone tissue engineering: in vitro and in vivo studies. Scientific Reports, 2020. 10(1): p. 1–14.

13. Nekounam, H., et al., Simple and robust fabrication and characterization of conductive carbonized nanofibers loaded with gold nanoparticles for bone tissue engineering applications. Materials Science and Engineering: C, 2020. 117: p. 111226.

14. De Santis, R., et al., Towards the design of 3D fiber-deposited poly (-caprolactone)/iron-doped hydroxyapatite nanocomposite magnetic scaffolds for bone regeneration. Journal of biomedical nanotechnology, 2015. 11(7): p. 1236–1246.

15. Kim, J.-J., et al., Magnetic scaffolds of polycaprolactone with functionalized magnetite nanoparticles: physicochemical, mechanical, and biological properties effective for bone regeneration. Rsc Advances, 2014. 4(33): p. 17325–17336.

16. Ji, L., et al., α-Fe2O3 nanoparticle-loaded carbon nanofibers as stable and high-capacity anodes for rechargeable lithium-ion batteries. ACS applied materials & interfaces, 2012. 4(5): p. 2672–2679.

17. Ji, L., et al., Electrospun carbon nanofibers decorated with various amounts of electrochemically-inert nickel nanoparticles for use as high-performance energy storage materials. Rsc Advances, 2012. 2(1): p. 192–198.

18. Ji, L., A.J. Medford, and X. Zhang, Porous carbon nanofibers loaded with manganese oxide particles: Formation mechanism and electrochemical performance as energy-storage materials. Journal of Materials Chemistry, 2009. 19(31): p. 5593–5601.

19. Kim, C., et al., Fabrication of electrospinning-derived carbon nanofiber webs for the anode material of lithium-ion secondary batteries. Advanced Functional Materials, 2006. 16(18): p. 2393–2397.

20. Toprakci, O., et al., Fabrication and electrochemical characteristics of electrospun LiFePO4/carbon composite fibers for lithium-ion batteries. Journal of Power Sources, 2011. 196(18): p. 7692–7699.

